# Monitoring alpha-synuclein oligomerization and aggregation using bimolecular fluorescence complementation assays: what you see is not always what you get

**DOI:** 10.1101/2020.05.02.074161

**Authors:** Bryan Frey, Abdelrahman AlOkda, Matthew. P. Jackson, Nathan Riguet, James A. Duce, Hilal A. Lashuel

**Author notes:** To whom correspondence should be addressed: Hilal A. Lashuel, Brain Mind Institute, Swiss Federal Institute of Technology Lausanne (EPFL), Lausanne, CH-1015, Switzerland.

## Abstract

Bimolecular fluorescence complementation (BiFC) was introduced a decade ago as a method to monitor alpha-synuclein (α-syn) oligomerization in intact cells. Since then, several α-syn BiFC cellular assays and animal models have been developed based on the assumption that an increase in the fluorescent signal correlates with increased α-syn oligomerization or aggregation. Despite the increasing use of these assays and models in mechanistic studies, target validation and drug screening, there have been no reports that 1) validate the extent to which the BiFC fluorescent signal correlates with α-syn oligomerization at the biochemical level; 2) provide a structural characterization of the oligomers and aggregates formed by the BiFC fragments; or 3) investigate the extent to which the oligomers of the fluorescent complex resemble oligomers formed on the pathway to α-syn fibrillization. To address this knowledge gap, we first analysed the expression level and oligomerization properties of the individual constituents of α-syn-Venus, one of the most commonly used BiFC systems, in HEK-293 & SH-SY5Y cells from three different laboratories using multiple approaches, including size exclusion chromatography, semiquantitative Western blot analysis, in-cell crosslinking, immunocytochemistry and sedimentation assays. Next, we investigated the biochemical and aggregation properties of α-syn upon co-expression of both BiFC fragments. Our results show that 1) the C-terminal-Venus fused to α-syn (α-syn-Vc) is present in much lower abundance than its counterpart with N-terminal-Venus fused to α-syn (Vn-α-syn) ; 2) Vn-α-syn exhibits a high propensity to form oligomers and higher-order aggregates; and 3) the expression of either or both fragments does not result in the formation of α-syn fibrils or cellular inclusions. Furthermore, our results suggest that only a small fraction of Vn-α-syn is involved in the formation of the fluorescent BiFC complex and that some of the fluorescent signal may arise from the association or entrapment of α-syn-Vc in Vn-α-syn aggregates. The fact that the N-terminal fragment exists predominantly in an aggregated state also indicates that one must exercise caution when using this system to investigate α-syn oligomerization in cells or *in vivo*. Altogether, our results suggest that cellular and animal models of oligomerization, aggregation and cell-to-cell transmission that are based on the α-syn BiFC systems should be thoroughly characterized at the biochemical level to ensure that they reproduce the process of interest and measure what they are intended to measure.

**Graphical Abstract:** 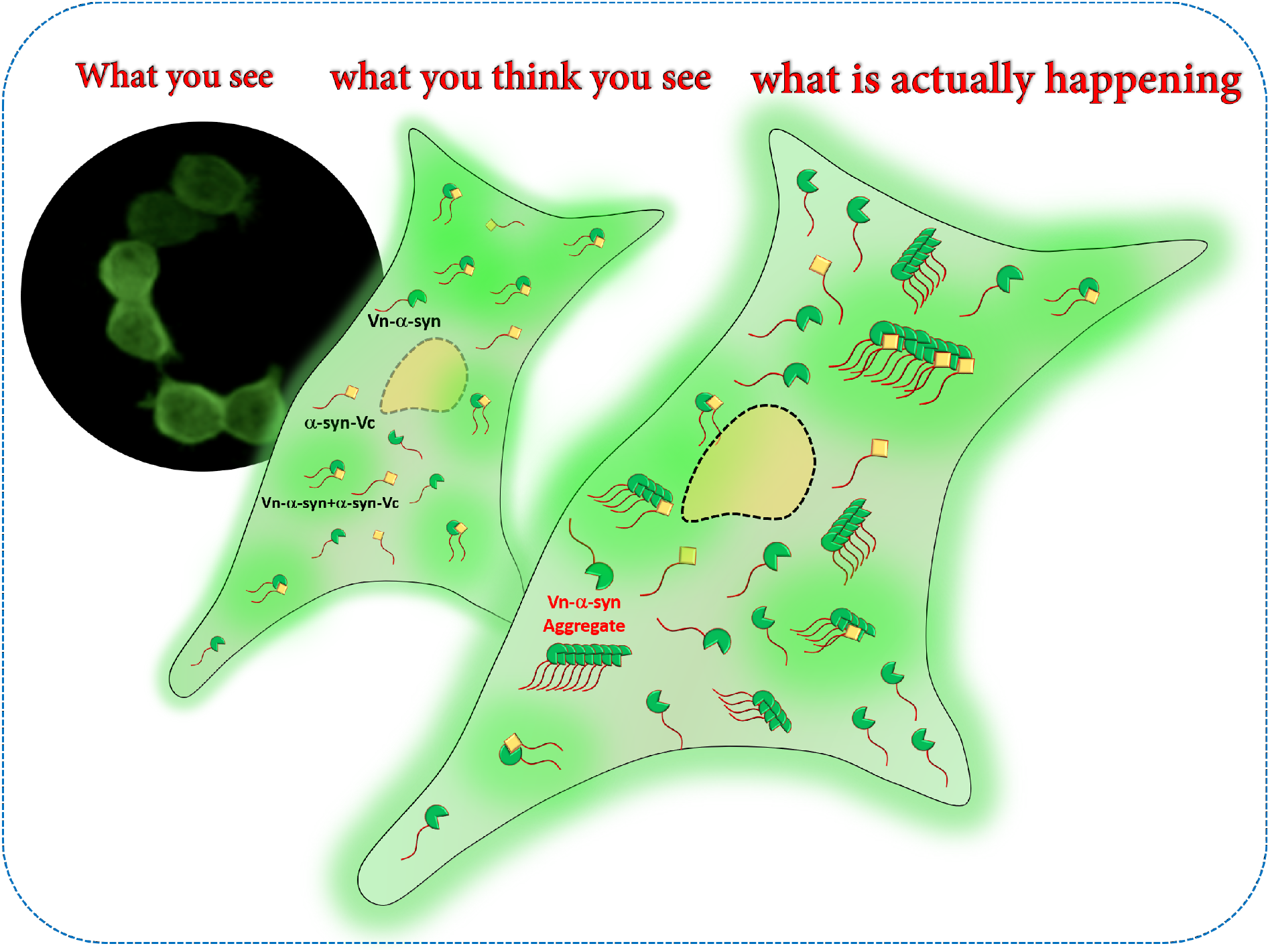

Bimolecular fluorescence complementation (BiFC) was introduced a decade ago to monitor alpha-synuclein oligomerization in intact cells, based on the assumption that an increase in the fluorescent signal correlates with α-synuclein oligomerization and aggregation. Herein, we used several biochemical and cellular assays to characterize commonly used α-synuclein Venus BiFC assays. Our results show that one of the BiFC fragments (Vn-α-synuclein) exhibits higher expression levels and aggregation propensity than its counterpart (α-synuclein-Vc), thus complicating the interpretation of the molecular interactions that give rise to the fluorescence signal and raise concerns about their application to investigate α-syn oligomerization in cells or *in vivo*.

## Introduction

α-synuclein (α-syn) is a soluble protein that is highly expressed in neurons of the central nervous system (Fauvet et al., 2012; Lashuel, Overk, Oueslati, & Masliah, 2012). It is a member of the synuclein family of proteins, which includes two other shorter proteins, namely, β-syn and γ-syn. Although the three proteins share significant sequence homology, only α-syn has been shown to aggregate and accumulate in a fibrillar form in Lewy bodies and Lewy Neurites; the pathological hallmarks of Parkinson’s disease (PD) as well as other neurodegenerative diseases including Alzheimer’s disease, dementia with Lewy bodies and multiple system atrophy (Goedert, Jakes, & Spillantini, 2017). Under physiological conditions, α-syn shuttles between different organelles and exists in multiple conformations depending on its localization and interactions with other proteins, lipids and membranes (Sulzer & Edwards, 2019). The transition of α-syn from a native state to its pathological forms requires it to undergo major conformational and quaternary structural changes involving a transition from disordered monomers to a mixture of secondary structured oligomers and then to β-sheet rich fibrillar aggregates (Fig. 2B). Several cellular and animal models that reproduce different aspects of α-syn aggregation and pathology formation have been developed (M.-B. Fares et al., 2016; Lee & Lee, 2002; Reyes et al., 2014; Volpicelli-daley et al., 2011). However, a lack of tools that enable the monitoring of early events associated with α-syn misfolding and oligomerization has limited the utility of these models in investigating the molecular and cellular determinants of α-syn oligomer formation and toxicity. Increasing evidence suggests that targeting early aggregation events and inhibiting α-syn oligomerization represents one of the most effective strategies for preventing α-syn pathology formation and treating PD. The first step towards achieving this therapeutic goal is the development of robust cellular and animal models in which such early oligomerization events could be visualized and quantified. Despite the reported development of several antibodies that specifically detect α-syn oligomers, but not monomers or fibrils, none have emerged as a validated oligomer-specific or robust tool for monitoring and quantifying α-syn oligomerization in cellular models (Kumar et al., 2020). Although several groups report on the development and application of different assays to quantify α-syn oligomers, the majority of these assays do not differentiate the aggregation states of α-syn (oligomers vs fibrils) or capture the diversity of α-syn oligomers and aggregates.

**Figure 1:**
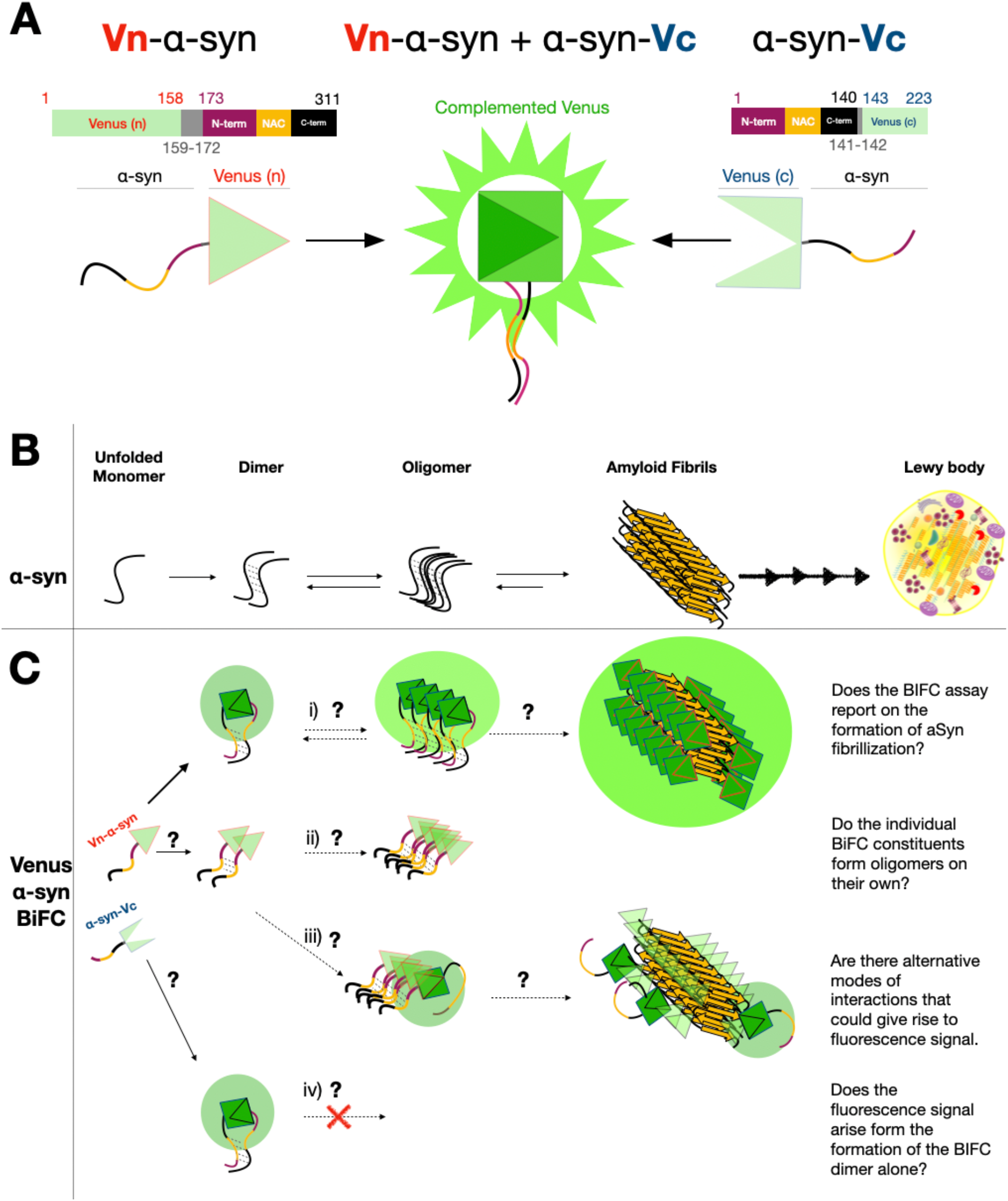
α-synuclein bimolecular fluorescence complementation (BiFC) assays: General principles and outstanding questions. **A.** Schematic depiction of the individual BiFC fragments which emits fluorescence upon α-syn-dimerization/oligomerization mediated reconstitution of the fluorescent protein Venus. **B.** A simplified representation of the mechanism of WT α-syn oligomerization, fibrillization and Lewy body formation **C**. Representation of the various possibilities for homo- and hetero-association between the BiFC fragments and that the scenarios that could lead to the generation of the BiFC fluorescence signal. From top to bottom: i) assumed α-syn BiFC oligomerization model, where the BIFC signal is generated upon α-syn-dimerization/oligomerization mediated reconstitution of the fluorescent protein, Venus. What is not known is whether this dimer is capable of forming higher order oligomers and/or convert to amyloid fibrils similar to α-syn fibrils found in Lewy Bodies. ii) Possibility that one or both of the two constituents self-oligomerize on their own. iii) illustration of how interactions between one monomeric BiFC constituent with an aggregated/oligomerized species of its counterpart could also give rise to fluorescence signals and thus wrongly interpreted as being derived from two monomers. iv) Reported drawback of BiFC: the formation of irreversible α-syn dimers with the reconstituted BiFC reporter that may not exhibit the same biophysical and oligomerization properties of native untagged α-syn. Vn-α-syn: (amino terminal fraction of Venus)-α-syn; α-syn-Vc: α-syn-(carboxy terminal fraction of Venus).

**Figure 2:**
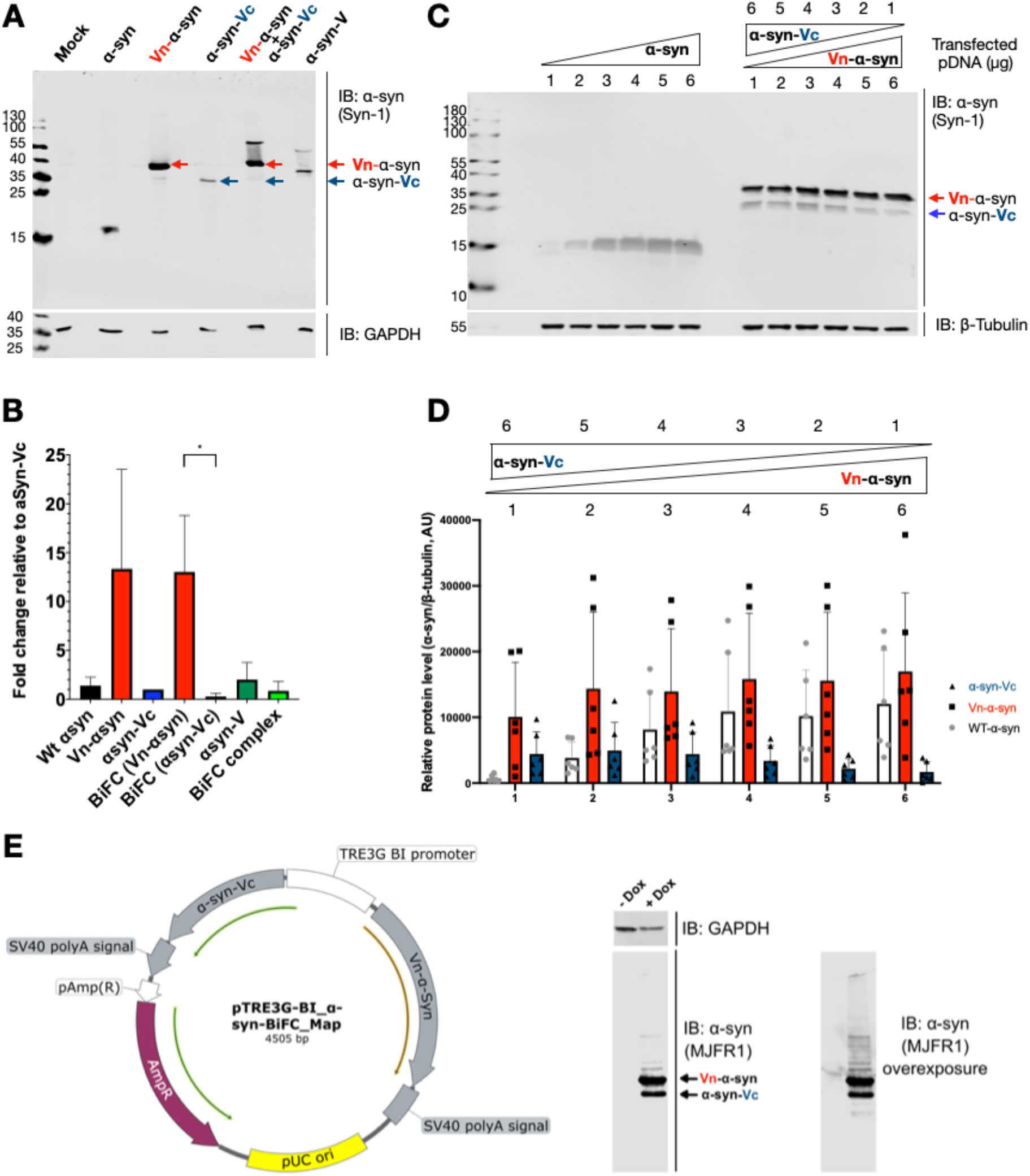
The two BiFC fragment proteins are not expressed at the same level in HEK-293 cells. **A.** HEK-293 cells overexpressing distinct α-syn constructs were lysed 48 h after transfection, and the cell lysates were analysed by immunoblotting for α-syn (Syn-1), without boiling. GAPDH was used as the loading control. A representative blot of n=3 experiments is shown **B.** Densitometry of data shown in (A.); a significant difference was indicated (p=0.045) by one-way ANOVA followed by a Tukey test for BiFC (Vn-α-syn) vs. BiFC (α-syn-Vc). **C.** WB of lysates from HEK-293 cells transfected with various amounts of vector α-syn DNA (depicted above the blot). α-syn was detected by anti-α-syn (Syn-1). The samples were boiled to disrupt complex formation to allow accurate determination of the levels of the two fragments. **D.** Relative protein levels (α-syn/β-tubulin) of BiFC proteins determined by quantification of the respective bands in (D) using Image Studio Lite. Data are shown as the mean of three independent experiments, each performed as technical duplicates (error bars represent the S.D.). **E.** Schematic of bidirectional inducible plasmid for expression of Vn-α-syn & α-syn-Vc under the control of a single tetracycline promoter. Immunoblot showing expression of both Vn-α-syn & α-syn-Vc upon addition of 1 μg /mL doxycycline. Samples were boiled prior to SDS-PAGE. No relevant significant difference was observed between the tested conditions using one-way ANOVA followed by a Tukey test. Vn-α-syn: (amino terminal fraction of Venus)-α-syn; α-syn-Vc: α-syn-(carboxy terminal fraction of Venus); α-syn-V (α-synuclein-Venus full length)

To allow monitoring of early oligomerization events (e.g. dimerization) involving α-syn in living cells, Outeiro *et al.* developed an α-syn bimolecular fluorescent complementation (BiFC) assay (Outeiro et al., 2008). The main principle behind this assay is the fusion of N- and C-terminal fragments of a reporter protein (e.g., GFP or YFP) to the proteins of interest, which in this case are α-syn monomers. These fragments are not fluorescent when they are separated and only emit a fluorescence signal if the two fragments are brought into close proximity to enable their association and the reconstitution of the structure of the full-length fluorescent protein; in this case, this occurs upon α-syn dimerization or oligomerization (Fig. 2A) (Kerppola, 2006, 2008). It is worth pointing out that BiFC was initially developed to study heterodimeric protein interactions (Hu et al., 2002; Hu & Kerppola, 2003), and its applications were later extended to investigate the self-assembly of amyloid-forming peptides and proteins (Outeiro et al., 2008).

The α-syn-BiFC system has been used to develop cellular assays to investigate several aspects of α-syn aggregation and biology, including 1) investigating the molecular and sequence determinants of α-syn oligomerization and aggregation (Aelvoet et al., 2014; Bae et al., 2014; K M Danzer et al., 2011; Karin M. Danzer et al., 2012; Delenclos et al., 2016; Gonçalves et al., 2016; Putcha et al., 2010; Savolainen et al., 2015; Tetzlaff et al., 2008; Zheng et al., 2018); 2) α-syn cell-to-cell transmission and propagation (Bae et al., 2014; Kim et al., 2016); 3) the screening of genetic (Dettmer et al., 2015; Lázaro et al., 2016, 2014) and pharmacological modifiers (Dominguez-Meijide et al., 2020; Mohammadiet al., 2017; Moussaud et al., 2015) of α-syn oligomerization, aggregation and clearance; and 4) the validation of therapeutic targets (Outeiro et al., 2008). This approach was also extended to animal models of synucleinopathies, such as rats (Dimant et al., 2013), *C. elegans* (Kim et al., 2016) and, most recently, *Drosophila* (Prasad et al., 2018) and mice (Cai et al., 2018) (Kiechle et al., 2019).

While reviewing the literature, we noted the lack of data on 1) the biochemical validation of this system, i.e., evidence that reports on the formation of α-syn dimers, oligomers or higher-order aggregates; 2) the stability and oligomerization properties of the BiFC fragments; 3) whether the fluorescent signal arises from an interaction between α-syn monomers only or between monomers of one fragment and oligomers or aggregates of the other fragment; and the biochemical, structural or morphological properties of the oligomers/aggregates formed upon co-expression of the BiFC fragments and to what extent they compare to α-syn aggregates found in diseased brains or formed during the fibrillization of recombinant α-syn *in vitro*. To address this knowledge gap, we performed a more in-depth characterization of the α-syn-Venus BiFC system in HEK-293 & SH-SY5Y cells that were transfected by different techniques and originated from three different laboratories using a combination of biochemical and cellular assays. Our results show that differential expression of the fluorescent protein fragments and the high propensity of one of these fragments to self-associate on their own complicate the interpretation of the fluorescence signal in this BiFC cellular assay. These observations highlight the limitations of using this system to investigate mechanisms and modifiers of α-syn oligomerization or aggregation. They also underscore the critical importance of validating BiFC-based assays at both a biochemical and ultrastructural level. This is crucial to determine whether the BiFC assays accurately reproduce and report on the process of interest, e.g., oligomerization, prior to their use in investigating mechanisms of protein oligomerization, amyloid formation or cell-to-cell transmission as well as screening modifiers of these processes.

## Materials and Methods

### Supporting Information

### Antibodies

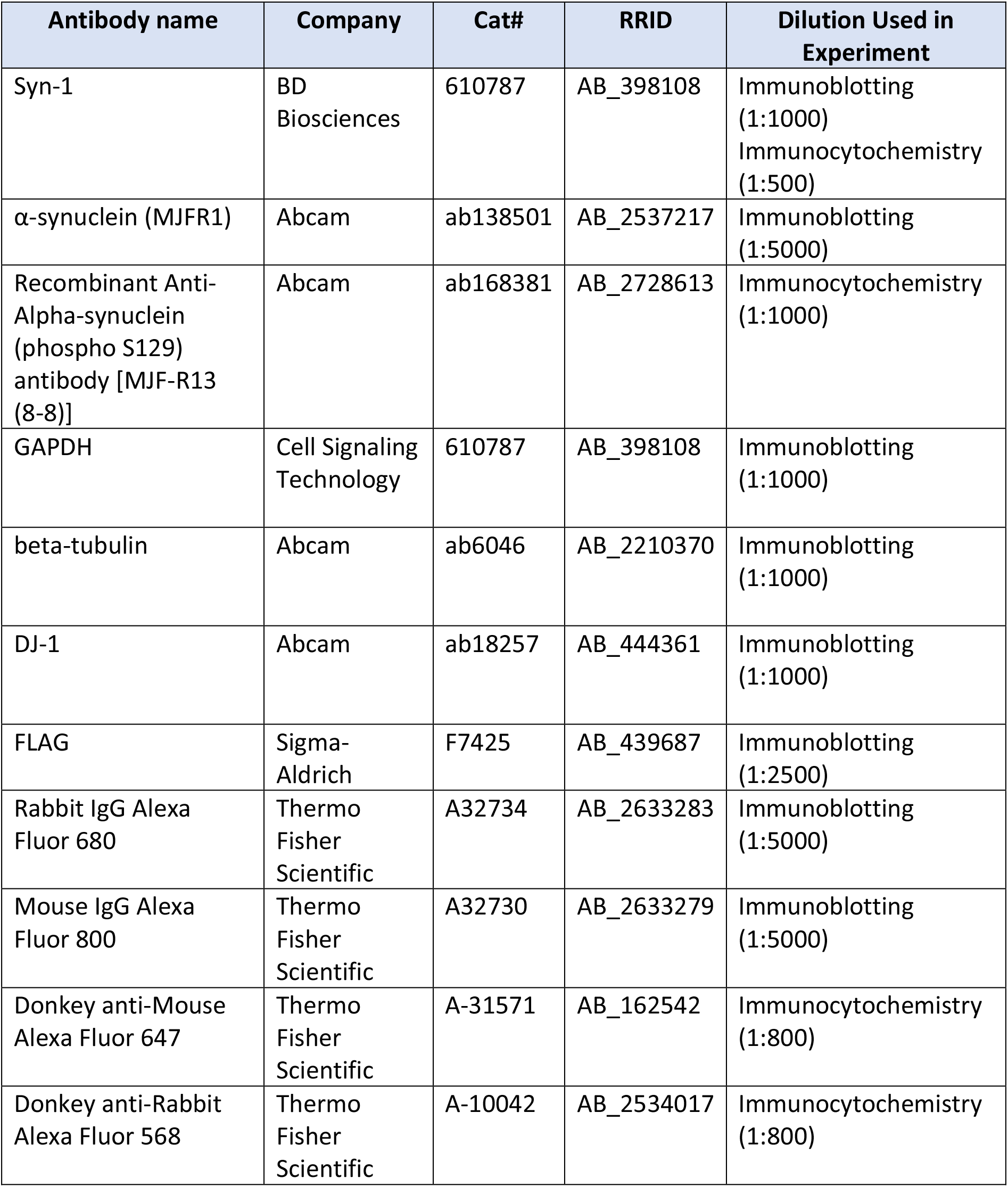

| Antibody name | Company | Cat# | RRID | Dilution Used in Experiment |
| --- | --- | --- | --- | --- |
| Syn-1 | BD Biosciences | 610787 | AB_398108 | Immunoblotting (1:1000)<br>Immunocytochemistry (1:500) |
| α-synuclein (MJFR1) | Abcam | ab138501 | AB_2537217 | Immunoblotting (1:5000) |
| Recombinant Anti-Alpha-synuclein (phospho S129) antibody [MJF-R13 (8-8)] | Abcam | ab168381 | AB_2728613 | Immunocytochemistry (1:1000) |
| GAPDH | Cell Signaling Technology | 610787 | AB_398108 | Immunoblotting (1:1000) |
| beta-tubulin | Abcam | ab6046 | AB_2210370 | Immunoblotting (1:1000) |
| DJ-1 | Abcam | ab18257 | AB_444361 | Immunoblotting (1:1000) |
| FLAG | Sigma-Aldrich | F7425 | AB_439687 | Immunoblotting (1:2500) |
| Rabbit IgG Alexa Fluor 680 | Thermo Fisher Scientific | A32734 | AB_2633283 | Immunoblotting (1:5000) |
| Mouse IgG Alexa Fluor 800 | Thermo Fisher Scientific | A32730 | AB_2633279 | Immunoblotting (1:5000) |
| Donkey anti-Mouse Alexa Fluor 647 | Thermo Fisher Scientific | A-31571 | AB_162542 | Immunocytochemistry (1:800) |
| Donkey anti-Rabbit Alexa Fluor 568 | Thermo Fisher Scientific | A-10042 | AB_2534017 | Immunocytochemistry (1:800) |

### Plasmids

The plasmid encoding WT-α-syn was cloned into a pAAV vector under the control of a CMV promoter (Addgene plasmid #36055; http://n2t.net/addgene:36055; RRID: Addgene36055). The Vn-α-syn (Addgene plasmid #89470; http://n2t.net/addgene:89470; RRID:Addgene_89470) and α-syn-Vc (Addgene plasmid #89471; http://n2t.net/addgene:89471; RRID:Addgene_89471) plasmid fragments were both previously cloned into the backbone of pcDNA3.1 under the control of a CMV promoter and were a kind gift of Tiago Outeiro. The α-syn-Venus (α-syn-V) construct encoding α-syn fused to full-length Venus under the control of a CMV promoter in a pcDNA 3.1 backbone was a kind gift from Pamela McLean. BacMam viral plasmids & production was performed by Oxford Expression Technologies (OET) using the template DNA from Addgene plasmids #89470 & #89471. Takara Clontech pTRE3g bidirectional plasmid (Takara Clontech Cat# 631337) for inducible expression was subcloned from Addgene plasmids #89470 & #89471. Further details about each vector and the full sequences can be obtained upon request.

### Cell culture and transfection protocol

HEK-293 cells were maintained in Dulbecco’s modified Eagle medium (DMEM; Gibco, Cat# 31966-021) containing 10% FBS and streptomycin and penicillin (10 μg/ml) at 37 °C. The maximum number of passages was 15 times. Cells were transfected by the addition of polyethylenimine (PEI) (Polysciences Inc, Cat# 23966) to plasmid DNA in Opti-MEM (Gibco, # 31985-062) at a ratio of 1:3 (μg vector DNA:μg PEI). The added transfection mixture volume was 10% of the total volume of the cell culture medium present in each well. The amount of plasmid DNA used for transfection was 2 μg for 6-well plates and 10 μg for 10 cm dishes. For BiFC co-expression (Vn-α-syn and α-syn-Vc), 1 μg of each BiFC vector was used for 6-well plates, and 5 μg of each vector was used for 10 cm dishes.

Induced expression of the bidirectional plasmid required the use of HEK 293 Tet-On^®^ 3G cells (Takara Clontech Cat# 631182). Cells in 6 well plates were transfected with 1 μg of the bidirectional plasmid using lipofectamine 3000 (Thermo Fisher Scientific Cat# L3000-008) as per manufacturer instructions. 24 h after transfection expression was induced with 1 μg / mL doxycycline.

For BacMam viral studies, SH-SY5Y (ECACC) neuroblastomas were maintained in 1:1 DMEM:F12 (Thermo Fisher Scientific Cat# 11-039-021) containing 10% FBS and streptomycin and penicillin (10 μg/ml) at 37 °C. The maximum number of passages was 20. BacMam virus was added at a multiplication of infectivity of 200 according to viral titre determined by OET. For BiFC expressing cell lines, SH-SY5Y (ECACC) neuroblastomas stably expressing either Vn-α-syn or α-syn-Vc were maintained in DMEM-Glutamax (Thermo Fisher Scientific Cat# 1056601610566016) containing 10% FBS, streptomycin and penicillin (10 μg/ml) and Geneticin (300μg/ml) at 37 °C. The maximum number of passages was 10. SH-SY5Y (ECACC) cell lines that either stably expressed Vn-α-syn or α-syn-Vc were a kind gift from Dr. Seung-Jae Lee (Bae et al., 2014). These cells were maintained in DMEM-Glutamax (Thermo Fisher Scientific Cat# 1056601610566016) containing 10% FBS, streptomycin and penicillin (10 μg/ml) and Geneticin (300μg/ml) at 37 °C. The maximum number of passages was 10.

### Preparation of α-syn fibrils

Pre-formed fibrils (PFFs) were generated, characterized as used in the neuronal seeding model as described previously in (Mahul-Mellier et al., 2020).

### Immunocytochemistry

HEK-293 cells (300 000 cells per well in 2 ml cell medium) were seeded onto coverslips coated with poly-L-lysine (Cultrex, Cat# 3438-100-01) and transfected as described above. Cells were washed 2 times with PBS and fixed in 3.7% formaldehyde (Sigma Aldrich) at RT for 15 min. Cells were then blocked and permeabilized simultaneously in PBS containing 3% BSA and 0.1% Triton X-100 (PBS-T) at RT for 30 min. α-syn was targeted by using the Syn-1 antibody (BD Biosciences) in PBS-T (diluted 1:500) and the α-syn pSer129 antibody MJF-R13 (Abcam, ab168381) at RT for2 h. Cells were then rinsed five times with PBS-T and incubated with a cocktail of donkey anti-mouse Alexa Fluor 647 (1:800), donkey anti-rabbit Alexa Fluor 568 (1:800) and DAPI (1:1500) in PBS-T at RT for 1 h. Cells were then washed 4 times in PBS-T and finally rinsed with pure water before being mounted in polyvinyl alcohol (PVA) mounting medium with DABCO (Sigma Aldrich, Cat# 10981). Images of cells were captured with a confocal laser scanning microscope using a 40x oil objective and Zen software (LSM 700, Zeiss).

As positive control for the analysis of α-syn inclusion formation, primary hippocampal neurons were treated with PFFs for 14 days to induce the formation of α-syn inclusions. In this model, the inclusions are pS129, ubiquitin, p62 and Thioflavin S positive, as reported previously (Mahul-Mellier et al., 2020). Thus, enabling us to validate the protocols used to assess the aggregation state and amyloid-like properties of α-syn in the BiFC cellular assays.

### Cell lysis

HEK-293 or SH-SY5Y cells were plated in 6-well plates (300 000 cells per well in 2 ml cell medium) and lysed in RIPA buffer (150 mM NaCl, 50 mM Tris pH 8.0, 1% NP-40, 0.5% deoxycholate, and 0.1% SDS) supplemented with 1% protease cocktail inhibitor (Sigma Aldrich, Cat# PB8340) and 1% phenylmethylsulfonyl fluoride (Axon Lab, Cat# A0999.0005) 48 h after transfection. The cell suspension was then gently mixed while being rotated at 4 °C for 30 min, and the cell debris was pelleted afterwards by centrifugation at 4 °C and 13 000 g for 20 min. The supernatant fraction was then used for immunoblotting.

For analysis of the SDS-insoluble fractions, the samples were centrifuged for 30 min at 13 000 g instead of 20 min, and the insoluble fraction (pellet) was resuspended in TBS/SDS buffer (150 mM NaCl, 50 mM Tris, pH 7.5, and 2% SDS) and sonicated using a probe for 15 s (1 s on pulse/1 s off pulse) at a 20% amplitude. The soluble and insoluble fractions were then processed as described in the immunoblotting section.

### Immunoblotting procedure

The total protein concentration for each sample was determined by the bicinchoninic acid assay (BCA, Thermo Scientific, Cat# 23225). Samples were next mixed (4:1) with 4x Laemmli buffer (2% sodium dodecyl sulfate (SDS), 20% glycerol, 0.05% bromophenol blue, 0.125 M Tris-HCl pH 6.8 and 5% β-mercaptoethanol). Samples were not boiled to maintain the stability of the protein interactions. Total cell lysates (10 μg) were separated with 15% SDS-PAGE (polyacrylamide gel electrophoresis) and transferred to a nitrocellulose membrane (Chemie Braunschweig, Cat# MSNC02030301) with a semidry system (BioRad, Cat# 1704150) at a constant amperage of 0.5 A and 25 V for 45 min. The membranes were then blocked with Odyssey Blocking buffer (Li-Cor, Cat# 927-40000) that was diluted 1:3 in PBS at 4 °C overnight. The membranes were then probed with the respective primary antibody (1:1000 diluted in PBS) on a shaking platform at RT for 2 h and then washed 3 times for 10 min in PBS-T (PBS, 0.1% Tween 20). The membranes were next incubated with the respective Alexa-conjugated secondary antibodies (goat anti-mouse IgG Alexa Fluor 800 or goat anti-rabbit IgG Alexa Fluor 680; 1:5000 in PBS) on a shaking platform while protected from light at RT for 1 h. The blots were finally washed 3 times in PBS-T and scanned by an Odyssey CLx Imaging System (Li-Cor Biosciences) at a wavelength of 700 or 800 nm.

For BacMam-transduced samples in SH-SY5Y cells the above protocol was followed with the following exceptions. For native gel experiments 15 μg of each sample were mixed with native sample buffer (BioRad, Cat # 1610738) and loaded on NuPAGE 4–12% Bis-Tris gels (Thermo Scientific, Cat# NP0335BOX) with Tris-Glycine running buffer (BioRad, Cat# 1610734). Total protein was detected on the gel using InstantBlue Protein Stain (Expedeon Cat# ISB1L-EXP-1L) before fluorescence was visualised using a GE Amersham Typhoon (GE Healthcare Lifesciences) with the GFP filter set. Protein was then transferred to PVDF (BioRad Cat# 1704157) using a BioRad TurboBlot. Membranes were blocked with 5% milk and incubated with primary antibody diluted in blocking buffer at 4 °C overnight. For non-native experiments 15 μg of each sample were mixed with 4 × XT sample buffer (BioRad, Cat #1610791), loaded on Criterion XT 4–12% Bis-Tris gels (BioRad, Cat# N3450124) with NuPAGE MES-SDS running buffer (Thermo Scientific, Cat# NP000202).

### Crosslinking of intact cultured cells

Cells cultured in 6-well plates (300 000 cells in 2 ml medium) were removed from the plate by enzymatic digestion via trypsinization 48 h post-transfection and collected by centrifugation of the cells at 300 g and RT for 3 min. Subsequently, the cells were resuspended in 600 μl of PBS containing protease inhibitors (Roche, Cat# 05892791001) and split into two 300 μl aliquots. One aliquot was treated with DSG (disuccinimidyl glutarate) crosslinker (1 mM) (Thermo Scientific, Cat# 20593) and the other served as a DMSO only negative control. DSG was stored at 4 °C with desiccant and prepared fresh directly before use by dissolving it in DMSO to make a 100 mM solution (100x stock solution). Crosslinking was performed under shaking conditions at 750 rpm for 30 min at 37 °C. The reaction was then quenched by adding Tris (50 mM, pH 7.6) and incubating at RT for 15 min. Next, the cells were lysed by sonication (amplitude of 40%, 2 x 15 s) and centrifuged at 20 000 g for 30 min at 4 °C. The supernatant was collected, and total protein concentration determined by BCA. The sample was then mixed 1:1 with NuPAGE LDS sample buffer (Thermo Scientific, Cat# NP0007) and boiled at 95 °C for 10 min. Twenty micrograms of each sample was run on NuPAGE 4–12% Bis-Tris and proteins were electroblotted onto PVDF membranes (Millipore, ISEQ00010), which were further processed as described in the immunoblotting section.

For BacMam-transduced samples in SH-SY5Y cells the above protocol was followed with the following exceptions. Samples were cross-linked in plate, mixed with BioRad XT sample buffer (BioRad Cat# 1610738) and resolved on BioRad Criterion 4-12 % precast gels (BioRad Cat# 3450125) before being transferred to PVDF (BioRad Cat# 1704157) using a BioRad TurboBlot.

### Size exclusion chromatography

Cells were plated in 10 cm dishes (2,2 × 10^6^ cells) and transfected as previously described. Cells were lysed 48 h post-transfection by nonionic detergent lysis buffer (20 mM Tris pH 7.4, 150 mM NaCl, 1 mM EDTA, 0.25% NP40, and 0.25% Triton X-100) supplemented with 1% protease cocktail inhibitor (Sigma Aldrich, Cat# PB340) and 1% phenylmethylsulfonyl fluoride (Axon Lab, Cat# A0999.0005) under gentle rotation for 30 min at 4 °C. Cell lysates were then cleared by centrifugation at 13 000 g for 30 min at 4 °C, and 300 μl of the collected supernatant was directly injected onto a Superdex 200 HR 10/300 column (GE Healthcare Life Science, Cat# 17517501). To avoid any aggregation artefacts due to freezing and thawing of the samples, the cell lysates were directly subjected to size exclusion chromatography using an ÄKTA purifier (GE Healthcare Life Science). The column was washed with three column volumes (75 ml) of Milli-Q and equilibrated with three column volumes of filtered PBS (20 mM sodium phosphate, 150 mM NaCl, pH 7.4) at a flow rate of 0.3 ml/min. An injection loop with a 500 μl volume was used to introduce the analytes and washed with 5 ml of buffer manually using a plastic syringe before each run. The flow rate for each run was set to 0.5 ml/min at 4 °C, and the collection volume was set to 500 μl. From each size fraction, a 40 μl aliquot was directly mixed with 10 μl of 4x Laemmli buffer (2% sodium dodecyl sulfate (SDS), 20% glycerol, 0.05% bromophenol blue, 0.125 M Tris-HCl, pH 6.8 and 5% β-mercaptoethanol) and each sample was then subjected to immunoblotting as described in the corresponding section. For calibration, the Superdex 200 column was calibrated using a gel filtration HMW calibration kit (GE Healthcare Life Science, Cat# 28403842) that included six reference proteins with known molecular weights (MWs) (ranging from 43-669 kDa). The generated standard curve allowed us to calculate the apparent MW of each eluted α-syn construct (Fig. 4C). Dextran blue 2000 produced a peak at 7,78 ml, which defined the void volume (Vo) and thereby the maximal size of the proteins that could be separated (Fig. 4B).

**Figure 3.**
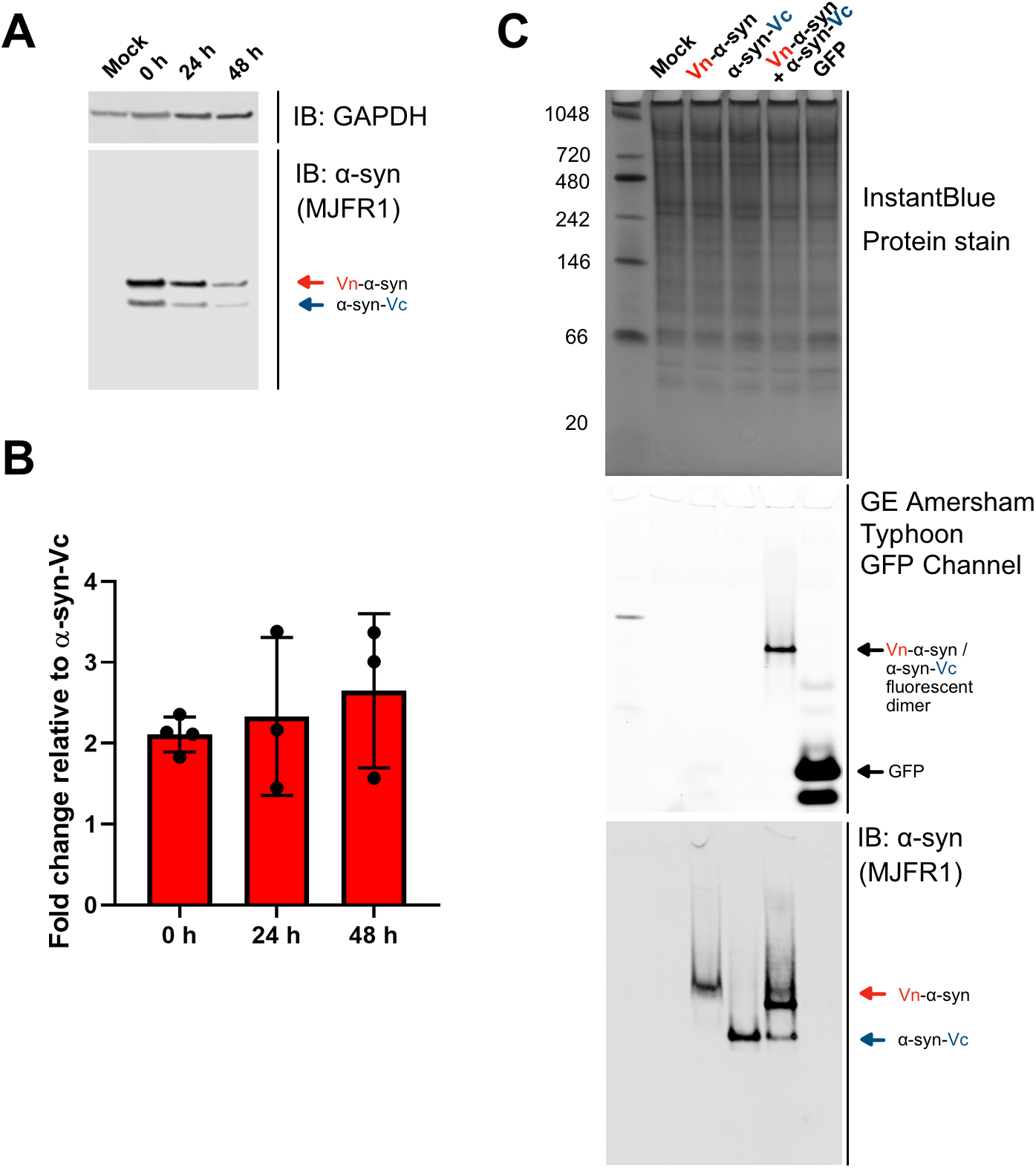
The two BiFC fragment proteins are not expressed to the same level in BacMam transduced SH-SY5Y cells. **A.** Denaturing immunoblot of Vn-α-syn & α-syn-Vc expression following BacMam virus mediated delivery in SH-SY5Y cells. Virus was added for 24 h, washed off and time 0 sample taken. Protein expression was tested 24 and 48 h after virus removal. **B.** Quantification of data displayed in A. using Image Studio Lite. Data are shown as the fold change Vn-α-syn normalised to α-syn-Vc (error bars represent the S.D.) at each time point. **C.** Native-PAGE analysis of detectable fluorescence following 24 h incubation with each respective BacMam virus. Samples were resolved by native-PAGE, equal protein loading confirmed with InstantBlue protein stain. In gel fluorescence imaged using a GE Amersham Typhoon Imager showing one main band for the Vn-α-syn & α-syn-Vc fluorescent dimer at a reduced intensity compared to BacMam-GFP virus. Immunoblot following transfer of native-PAGE to PVDF showing only small amount of Vn-α-syn & α-syn-Vc present as discrete dimer.

**Figure 4:**
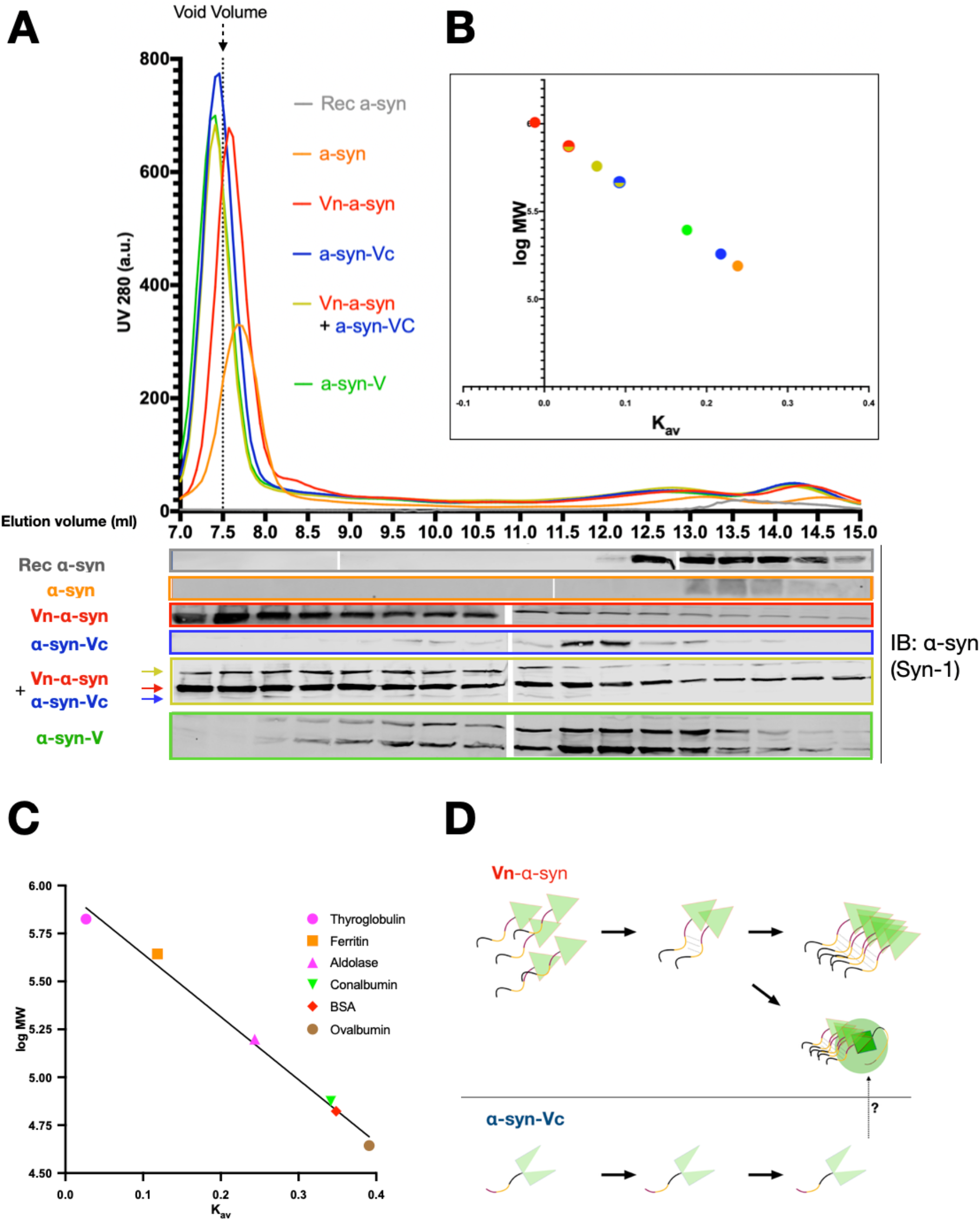
The Vn-α-syn fragment exhibits a high propensity to form soluble oligomers compared to α-syn-Vc and WT α-syn. **A.** Size exclusion chromatography of a HEK-293 cell lysate 48 h post-transfection with the indicated α-syn constructs. The SEC fractions were analyzed by WB using anti-α-syn (Syn-1) antibodies. **B.** Calculated apparent MW depicted with a log scale plotted against the Kav value of each α-syn construct (large colored dots). Mixed circles represent either Vn-α-syn (red) or α-syn-Vc (blue) when being co-expressed. **C.** Standard curve generated using the indicated six reference proteins. **D.** Working model suggesting that the high propensity of Vn-α-syn to oligomerize and how such oligomers could interact with or entrap α-syn-Vc. Note, the UV280 signal in A does not correspond to the α-syn signal, but mostly proteins in the remaining cell lysate. Representative data shown from n=3 experiments. Vo: dead volume; Ve: elution volume; Vn-α-syn: (amino terminal fraction of Venus)-α-syn; α-syn-Vc: α-syn-(carboxy terminal fraction of Venus); α-syn-V (α-synuclein-Venus full length).

### Statistical Analysis

The level of the corresponding α-syn constructs were estimated by measuring the WB band intensity using Image J software (U.S. National Institutes of Health, Maryland, USA; RRID:SCR_001935) and normalized to the relative protein levels of actin or GAPDH. The statistical analyses were performed using the ANOVA test followed by a Tukey-Kramer post-hoc test using Graph Pad Prism v.8.4.2. The data were regarded as statistically significant at p<0.05.

## Results

### The two α-syn-Venus BiFC fragments were expressed at different levels

As a first step in characterizing the Venus-α-syn BiFC system, we sought to assess the protein levels and stability of full-length α-syn either fused to the individual fragments or full-length Venus (α-syn-Venus). The constructs corresponding to these proteins were overexpressed in HEK-293 cells, and the cell lysates were analysed by Western blotting (WB) with antibodies against α-syn as well as the N- and C-terminal fragments of Venus (abbreviated as Vn-α-syn and α-syn-Vc, respectively). Both the full-length α-syn-Venus and Vn-α-syn protein levels were high, whereas the band corresponding to α-syn-Vc was barely detectable in comparison to that of Vn-α-syn (Fig. 2A). When expressed individually, Vn-α-syn was expressed at 6- to 16-fold higher levels than α-syn-Vc (Fig. 2A) and at levels higher than those of any of the other tested constructs (WT-α-syn, α-syn-Vc and α-syn-Venus), suggesting an increased stability or decreased turnover of Vn-α-syn that is consistent with previous observations (Dominguez-Meijide et al., 2020; Eckermann, Kügler, & Bähr, 2015; Gustafsson et al., 2018; Lázaro et al., 2014).

When the two constructs were co-expressed, the α-syn-Vc protein band became undetectable, while a new band corresponding to the Vn-α-syn + α-syn-Vc dimer was detected. This is consistent with previous reports of the irreversible dimerization of proteins upon successful reconstitution by BiFC (Hu et al., 2002). However, the band intensity of Vn-α-syn remained at a level that is comparable to that observed when the protein was expressed alone. These results suggest that only a small fraction of Vn-α-syn associates with α-syn-Vc that is involved in the formation of the fluorescent BiFC complex.

To assess whether it would be possible to increase the levels of α-syn-Vc and achieve an equimolar concentration of both BiFC constituents, we varied the amounts of vector DNA applied for transfection. As expected, the protein level of WT-α-syn was increased with the greater amount of vector DNA used for transfection (Fig. 2C). But, strikingly, despite the transfection of six times more α-syn-Vc than Vn-α-syn, the levels of α-syn-Vc remained unchanged (Fig. 2C-D). In parallel, we created a bidirectional expression system (pTRE3G-Bi Takara Clontech) that allowed for the inducible expression of both BiFC subunits fused with α-syn under the control of a single promoter (Fig. 2E). Upon transient transfection of this plasmid into HEK293 Tet-On cells (Takara Clontech) expression of both α-syn-Vc & Vn-α-syn was induced with 1 μg/mL doxycycline. Despite controlling expression in this manner an imbalance of protein expression was still observed with Vn-α-syn still being the dominant species (Fig. 2E). Surprisingly, this imbalance has been repeatedly observed but not discussed in previous studies (Cai et al., 2018; Dominguez-Meijide et al., 2020; Eckermann et al., 2015; Gustafsson et al., 2018; Lázaro et al., 2016, 2014; Prasad et al., 2018; Zondler et al., 2014). Only one report (Eckermann et al., 2015) suggested that the differential expression of the two constructs could lead to a potential entrapment of α-syn-Vc by the strongly overexpressed Vn-α-syn and force the less abundantly expressed protein into an non-specific interaction.

These data were consistent in orthogonal experiments including BacMam viral delivery of Vn-α-syn & α-syn-Vc into SH-SY5Y cells (Fig. 3). Whilst the α-syn-Vc protein band in this system was not reduced to a level that was undetectable, there was a consistently higher expression of Vn-α-syn over α-syn-VC across multiple time points ranging from 24 h to 72 h after exposure to virus (Fig. 3B). Venus fluorescence of BacMam delivered proteins resolved on a native-PAGE gel primarily revealed a small proportion of Vn-α-syn and α-syn-Vc dimer (compared to BacMam GFP) (Fig. 3C). Transfer of these proteins to a membrane and subsequent probing for α-syn by WB showed that only small amounts of the total α-syn correlated with the Vn-α-syn and α-syn-Vc dimer band that was fluorescent. This confirmed that only a small proportion of Vn-α-syn and α-syn-Vc dimerised into a form detectable by Venus fluorescence.

### Vn-α-syn exhibits a high propensity to form oligomers and higher-order aggregates

Next, we sought to investigate the biochemical and oligomerization properties of each of the BiFC fragments. Towards this end, the lysates of HEK-293 cells expressing the various α-syn BiFC proteins were separated by size exclusion chromatography (SEC). To assess whether any of the reagents or lysis of the cells affected size separation, the elution pattern of recombinant α-syn (Fig. 4B, grey line) was compared to that of the lysate of WT-α-syn-expressing HEK-293 cells (Fig. 4B, orange line) via analysis of the eluates by WB. Note, the resulting UV chromatogram of the cell lysates mostly represented total proteins rather than α-syn, which is why WB analysis was crucial for the comparison of the elution pattern. The elution of α-syn at 13-14 ml for both samples suggested that intracellular α-syn exists predominantly as a monomer, which is consistent with previous findings (Fauvet et al., 2012; Weinreb, Zhen, Poon, Conway, & Lansbury, 1996).

The largest α-syn construct, α-syn-Venus, eluted from 8 to 14,5 ml with the most intense band detected at 11-13 ml (Fig. 4B, green). This suggests that α-syn-Venus has a low propensity to form higher-order oligomers but that the majority of the protein exists in a monomeric form. Given that Vn-α-syn and α-syn-Vc have sizes that are between those of α-syn-V and WT-α-syn, we expected these proteins to elute at volumes between 8 ml and ~14 ml. As expected, α-syn-Vc eluted at 13 ml (Fig. 4B, blue) but in contrast the majority of Vn-α-syn eluted at the void volume of 7,5 ml (Fig. 4B, red), indicative of the high propensity for this protein to self-oligomerize. When both BiFC constructs were the band corresponding to the size of Vn-α-syn still eluted mainly at the void volume (Fig. 4B, yellow). Interestingly, co-expressed α-syn-Vc was now detected in higher MW fractions (<10 ml), possibly due to a shift caused by the dimerization of the BiFC constructs and consistent with what we observed by WB (Fig. 2A). Furthermore, the detection of a band in the void volume corresponding to the BiFC-α-syn dimer complex suggested that this complex goes on to form higher-order oligomers or that it is sequestered by the aggregates formed by the Vn-α-syn fragment (Eckermann et al., 2015). Additionally, we also analysed the lysate of SH-SY5Y cells that stably expressed either the α-syn-Vc or Vn-α-syn constructs using SEC, which confirmed an intrinsically high aggregation propensity of Vn-α-syn in independent system (Supplementary Figure 1).

### Crosslinking studies confirm that Vn-α-syn but not α-syn-Vc forms higher MW weight oligomers

To further characterize the aggregation propensities of the various α-syn BiFC fragments, we performed intact cell crosslinking and assessed the oligomerization by WB. DJ-1, which forms dimers in physiological conditions (Dettmer et al., 2015), was used as a positive control. Successful crosslinking within our experimental parameters was confirmed by the DSG-crosslinked cell lysate displaying a DJ-1 dimer band that was absent in non-crosslinked cells (Fig. 5B). WT α-syn was mainly detected as a monomer (15 kDa), but a noticeable stochastic ladder of dimers, trimers and higher molecular weight (MW) species was also observed (Fig. 5A). α-syn-Venus displayed two bands: one at the expected monomeric size of 42 kDa and one band at a lower size of approximately 36 kDa. The lower band was observed whenever the sample was not boiled prior to immunoblotting and is likely to represent a more compact folded structure of this fusion protein. In concordance with the SEC results, α-syn-Vc showed a single band at the expected MW, and no higher MW species were detectable. In contrast, Vn-α-syn formed several higher MW bands, including a prominent band that remained in the stacking gel. These high MW aggregates were only observed when intact cell crosslinking was performed. This suggests that in the absence of crosslinking, the majority of the Vn-α-syn oligomers are SDS-sensitive and break down into monomers in denaturing conditions. Additionally, the stacking gel showed a high-molecular-weight signal for α-syn as a result of co-expression, despite the intensity of this band being weaker than when Vn-α-syn alone was expressed. The aggregation seen here is likely to be driven by Vn-α-syn, as α-syn-Vc alone did not produce any higher MW aggregates. DSG crosslinking experiments were also performed in SH-SY5Y cells with BacMam virus mediated protein delivery (Fig. 5C). These data confirmed the differing oligomerization properties of the Vn-α-syn & α-syn-Vc proteins. Furthermore, immunodetection of a FLAG epitope present only in the α-syn-Vc construct confirmed that α-syn-Vc exists primarily as a monomer when crosslinked in the absence or presence of Vn-α-syn. Together, the results from the crosslinking and SEC experiments further establish that Vn-α-syn exhibits a high propensity to self-oligomerize and exists predominantly in oligomeric forms independent of α-syn-Vc.

**Figure 5:**
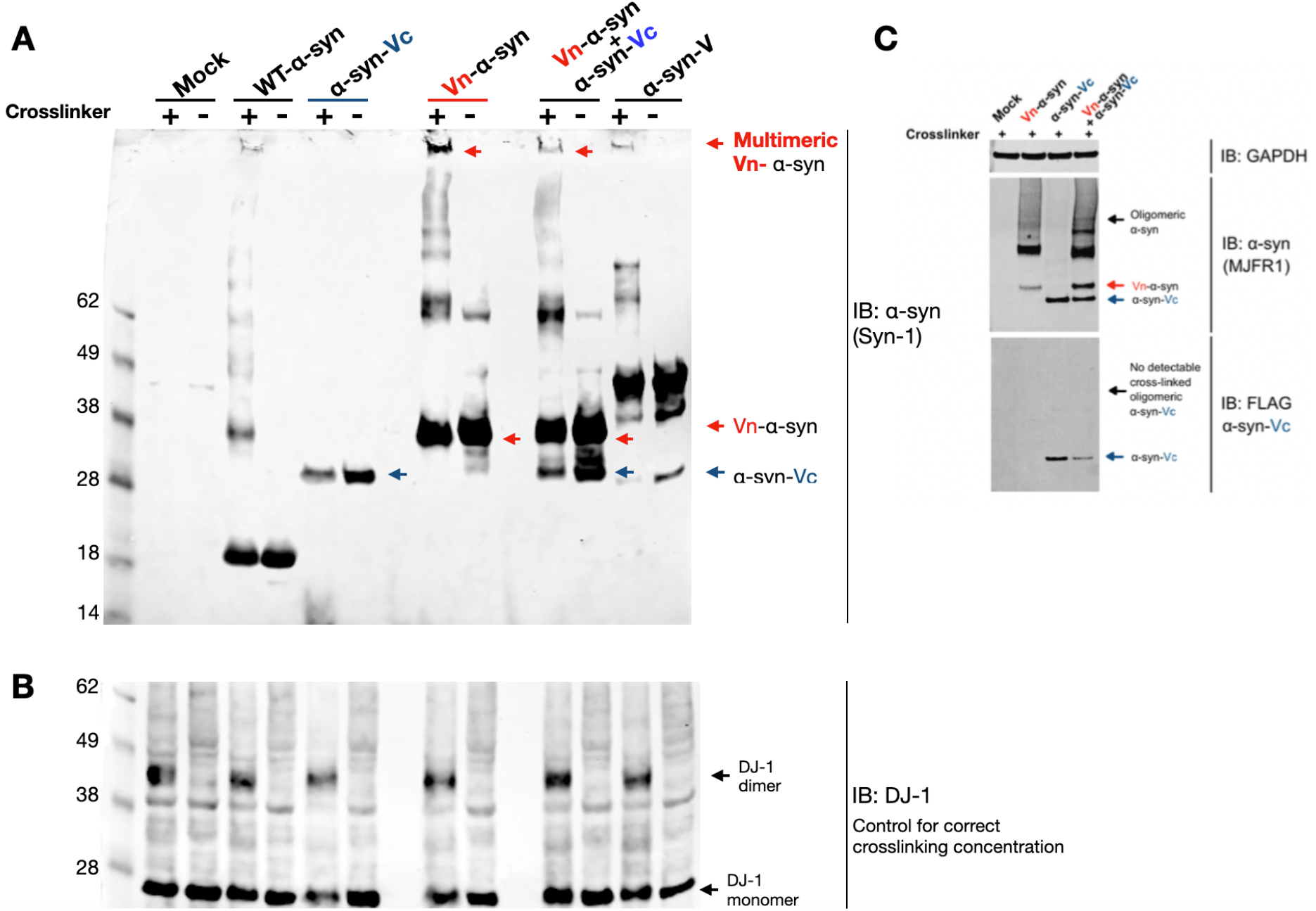
Intact cell crosslinking of α-syn BiFC constructs confirms SEC findings of Vn-α-syn oligomerization. **A.** Living HEK-293 cells were crosslinked by the addition of DSG 48 h post transfection with the indicated α-syn constructs, and the resulting cell lysate was analysed by Western blotting against α-syn using the Syn-1 antibody. All analysed α-syn proteins displayed higher MWs upon addition of the DSG crosslinker, except for α-syn-Vc. **B.** The physiological dimer DJ-1 was used as an internal control so that the correct crosslinker concentration was applied. DJ-1 immunoblotting confirms that dimers are only detected upon crosslinking. **C.** Immunoblot of DSG crosslinked to live SH-SY5Y cells following 24 h BacMam delivery of either Vn-α-syn or α-Syn-Vc or both constructs. All displayed samples treated with DSG. Cells treated with only Vn-α-syn displayed higher order crosslinked oligomeric species in addition to monomeric Vn-α-syn. Cells treated with only α-syn-Vc displayed only monomeric α-syn-Vc. Treatment with both Vn-α-syn and α-Syn-Vc displayed higher order crosslinked oligomeric species in addition to both monomeric proteins. To determine if the higher order crosslinked oligomeric species in the sample treated with both Vn-α-syn and α-Syn-Vc were comprised of both proteins a separate WB for the FLAG epitope present only in the α-Syn-Vc protein was performed, this revealed the higher order crosslinked oligomeric species were predominantly comprised of Vn-α-syn. EV: empty vector; Vn-α-syn: (amino terminal fraction of Venus)-α-syn; α-syn-Vc: α-syn-(carboxy terminal fraction of Venus); α-syn-V (α-synuclein-Venus full length). Blots are representative of n=3 experiments.

### Vn-α-syn predominantly forms soluble SDS-sensitive oligomers in cells

To determine whether the BiFC fragments form oligomers or higher-order aggregates such as fibrils, the lysates from HEK-293 cells expressing the different proteins were separated by centrifugation (at 13000g for 20-30 minutes) into soluble and insoluble fractions (Fig. 6A). This approach has been applied in previous studies to separate oligomeric and monomeric α-syn from insoluble α-syn species) (M.-B. Fares et al., 2016; Lee, Khoshaghideh, Patel, & Lee, 2004; Savolainen et al., 2015). WB analyses showed that Vn-α-syn was predominantly detected in the soluble fraction (Fig. 6B). This suggests that the self-assembled soluble oligomers of Vn-α-syn that eluted in the void volume peak (7,5 ml) of the SEC column (Fig. 4B) are disassociated into monomers when in the presence of SDS (Fig. 6B). In the insoluble fraction, the Vn-α-syn fragment was also detected as monomer band, suggesting that a small proportion of this fragment goes on to form insoluble aggregates. In contrast, none of the other constructs, including α-syn-Vc, could be detected in the insoluble fractions. Consistent with the data presented above, the ratio of insoluble over soluble fraction was higher for Vn-α-syn than for α-syn-Vc (Fig. 6C). Interestingly, when the two BiFC fragments were co-expressed, only Vn-α-syn (identified by size) was sufficiently enriched in the insoluble fraction. Only traces of α-syn-Vc and the complexed BiFC could be detected in the insoluble fraction, this could be due to the fact that these species become entrapped within or non-specifically interact with the Vn-α-syn aggregates. Similar observations were found in SH-SY5Y cells stably expressing either the Vn-α-syn or α-syn-VC (Supplementary Fig. 2).

**Figure 6:**
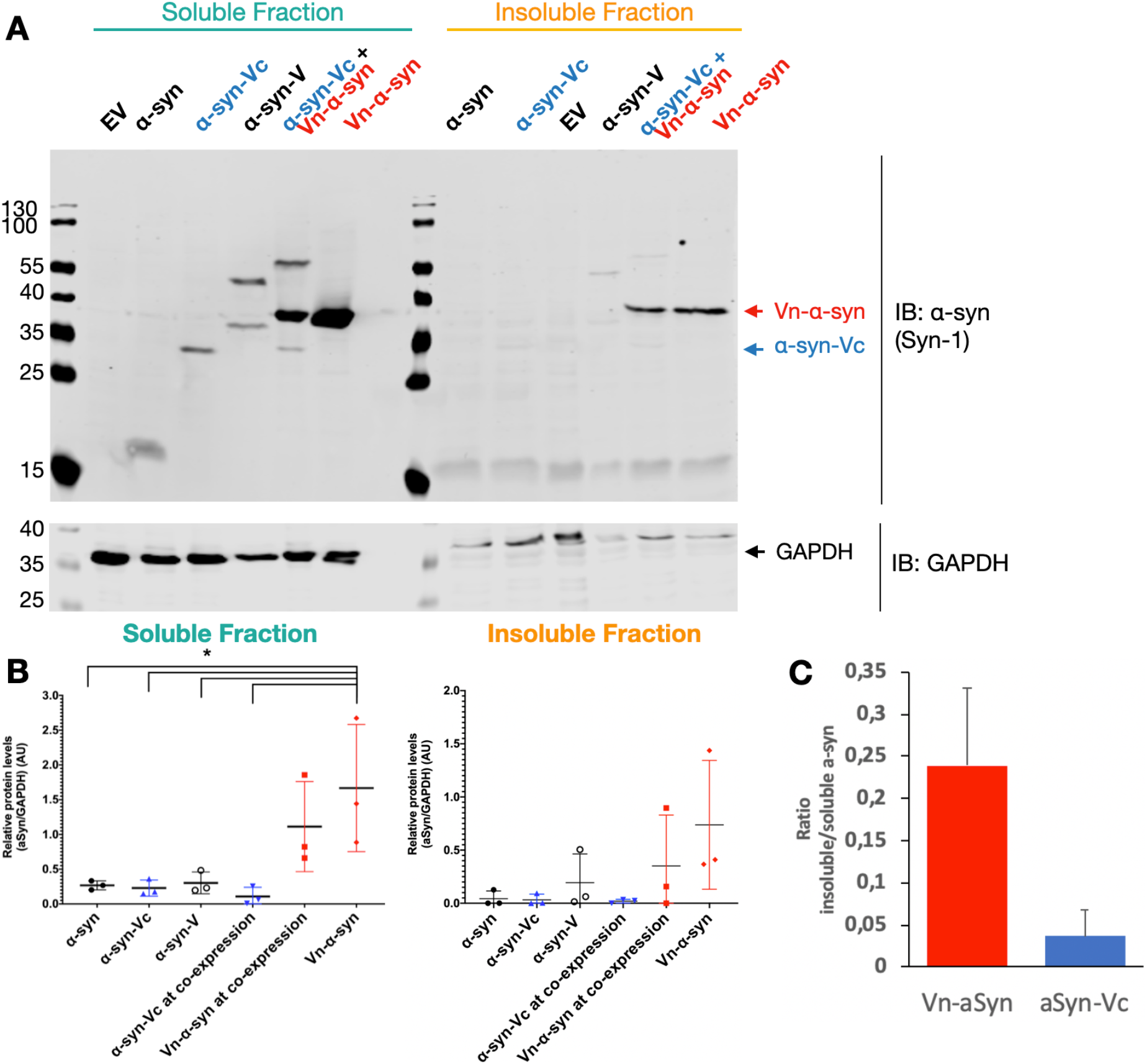
Vn-α-syn, but not α-syn-Vc or WT α-syn, is detected in the insoluble fractions. **A.** HEK-293 cells overexpressing distinct α-syn constructs were lysed 48 h after transfection, and the cell lysates were separated into a soluble and insoluble fraction by centrifugation. The resulting fractions were analysed by immunoblotting of α-syn (Syn-1). GAPDH was used as the loading control. Shown is a representative blot of n=3 experiments. **B.** Resulting soluble and insoluble band intensities of were quantified and normalized to soluble GAPDH. Significant differences were indicated (p<0.005) by one-way ANOVA followed by a Tukey test. **C.** Ratio of insoluble to soluble α-syn of Vn-α-syn and α-syn. Vc EV: empty vector; Vn-α-syn: (amino terminal fraction of Venus)-α-syn; α-syn-Vc: α-syn-(carboxy terminal fraction of Venus); α-syn-V (α-synuclein-Venus full length).

### Neither of the BiFC fragments when expressed alone or when co-expressed in cells form α-synuclein amyloid-like aggregates or inclusions

Finally, we sought to characterize the aggregation properties of the BiFC fragments in intact cells by immunocytochemistry. The proteins were expressed in HEK-293 cells, and their localization and aggregation were assessed using an α-syn specific antibody (Syn-1. α-syn fused to full-length Venus was used as a positive control and untagged α-syn as a negative control. As expected, no Venus fluorescence was observed when the individual BiFC fragments, Vn-α-syn or α-syn-Vc, were expressed alone, while co-expression of both proteins led to intense signals from Venus as a result of complementation (Fig. 7). Fluorescence was observed both in the cytosol and the nucleus, as previously reported (Dettmer et al., 2015; Lázaro et al., 2016, 2014; Moussaud et al., 2015). Consistent with the results from the biochemical and SEC studies, we did not observe the formation of foci or pSer129 positive inclusions for any of the BiFC constructs used, indicating that the BiFC fragments either alone or when co-expressed predominantly form soluble oligomers that do not exhibit amyloid-like properties, as evidence by the lack of binding to the amyloid-specific dye Amytracker (Supplementary Fig. 3). As a positive control, we used a primary neuronal α-syn seeding model where abundant α-syn fibrils are formed 7-14 days after treatment with preformed fibrils (PFFs). The α-syn aggregates in this neuronal model are both pS129 and Amytracker positive, consistent with previous reports by our groups and others (Mahul-Mellier et al., 2020). Although previous studies did not report quenching of the Amytracker signal by a GFP tag (Frottin et al., 2019; Rallis et al., 2020; Marrone *et al.,* 2020), we could not exclude that the presence of the Venus tag fused to α-syn could quench the fluorescence of Amytracker.

**Figure 7:**
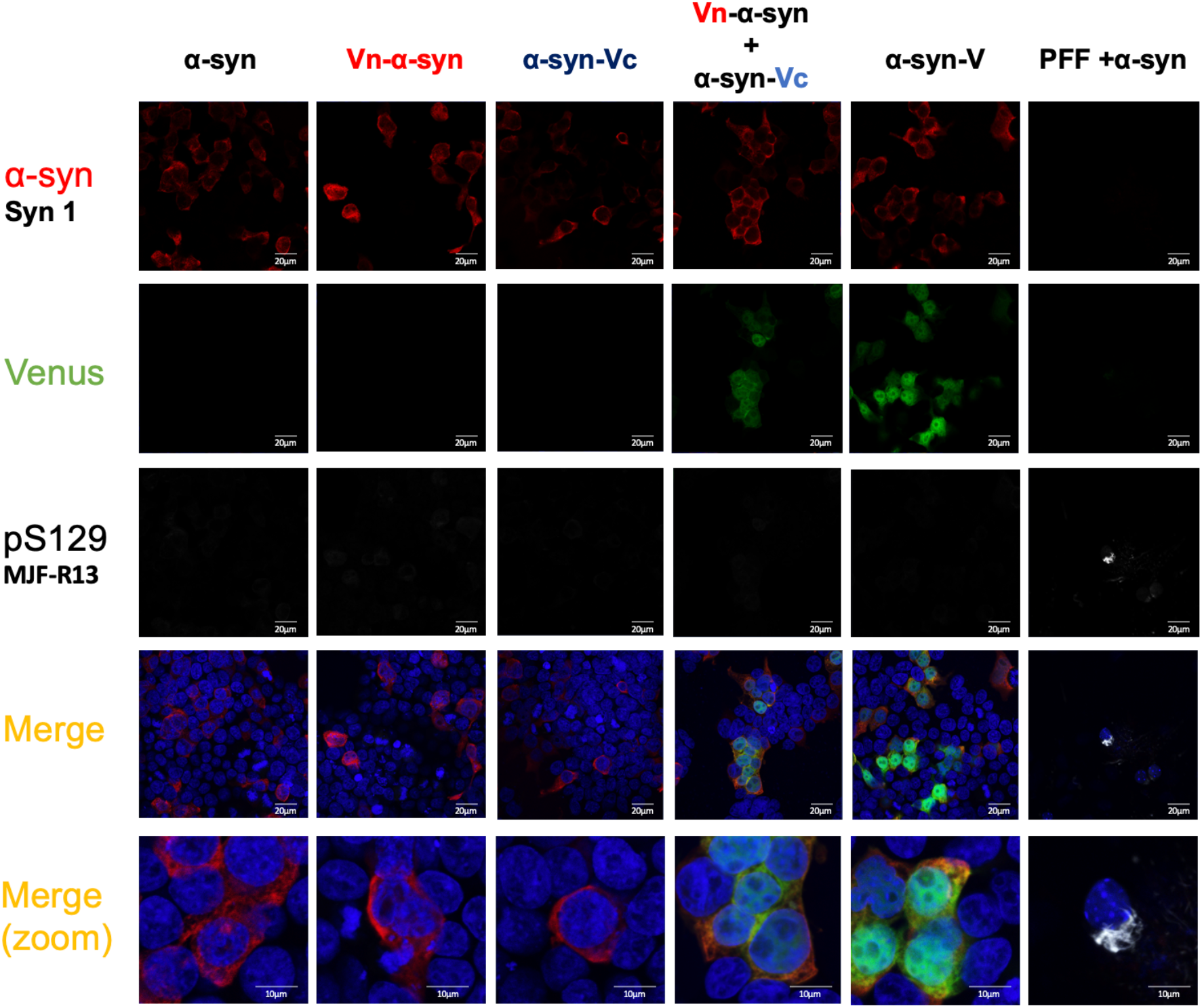
No α-syn inclusion, pSer129 signal or amyloid-like aggregates were observed in cell lines expressing the BiFC fragments. HEK-293 cells expressing the indicated α-syn constructs were fixed 48 h post-transfection and stained with the Syn-1 antibody. Upon co-transfection of the Vn-α-syn and α-syn-Vc constructs, Venus was successfully reconstituted and emitted a fluorescent signal that colocalized with the signal for α-syn. Mouse primary neurons were treated by PFF α-syn for 14 days and are positive for pS129. The nucleus was counterstained with DAPI. The scale bar is 20 μm in all the non-zoomed images and 10 μm for the zoomed images. The microscopy settings were kept the same among each construct. Shown are representative images from three independent experiments.

These observations demonstrate an absence of insoluble aggregated or fibrillar forms of α-syn in cells expressing either or both of the BiFC fragments, suggesting that the oligomers formed by the individual or complexed Venus BiFC fragments do not go on to form fibrils and may not share properties with oligomers that are known to be part of the pathway to α-syn fibril formation. Further studies are needed to characterize the biochemical and structural properties of the α-syn oligomers formed by the BiFC fragments to determine the extent to which they resemble native α-syn oligomers.

## Discussion

Several α-syn BiFC cellular assays (Gonçalves et al., 2010) have been developed and are increasingly used in cellular and animal studies aimed at identifying and validating the targets or mechanisms of α-syn aggregation, propagation and toxicity. These assays have also been used for screening genetic, enzymatic and pharmacological modulators of α-syn aggregation, toxicity and clearance (Cai et al., 2018; Dimant et al., 2013; Eastwood, Baker, Brooker, Frank, & Mulvihill, 2017; Kiechle et al., 2019; Moussaud et al., 2015; Outeiro, Putcha, Tetzlaff, Spoelgen, Koker, Carvalho, et al., 2008; Prasad et al., 2018). Despite this, there has been very little effort to validate these assays (Eckermann et al., 2015) at the biochemical level and to characterize the nature of the α-syn oligomers that form upon reconstitution of the α-syn BiFC fragments and fluorescence signal. The primary objective of this study was to contribute in addressing this knowledge gap by 1) providing insight into the molecular interactions that give rise to the fluorescence signal observed in these assays; and 2) investigate the solubility and aggregation state of the α-syn BiFC fragments and complexes.

Our studies show that α-syn-Vc is expressed at levels that are significantly less than Vn-α-syn in various cellular systems using different methods of protein delivery, and that only a fraction of Vn-α-syn participates in the formation of complexes that give rise to the fluorescence signal. This discrepancy in protein expression levels between the two fragments has been consistently observed in previous studies (Bartels et al., 2019; Cai et al., 2018; Delenclos et al., 2019; Eckermann et al., 2015; Gonçalves et al., 2016; Gustafsson et al., 2018; Herrera & Outeiro, 2012; Lázaro et al., 2016, 2014; Moussaud et al., 2015; Prasad et al., 2018; Zondler et al., 2014), but rarely discussed or taken into account in the interpretation of the results with the BiFC assays. Despite our attempts to express both proteins under a single Tet-On inducible promoter we could not overcome these differential expression issues. Interestingly, even when the Venus-BiFC fragments were expressed in Drosophila, protein levels of the α-syn-Vc fragment were significantly lower compared to the Vn-α-syn fragment (Prasad et al., 2018). Despite this observation the authors in this study chose to quantify the levels of insoluble α-syn using a filter retardation assay, a procedure that does not allow for the nature and distribution of the α-syn species in the insoluble fractions to be determined (i.e. the aggregates may consist purely of the aggregation-prone fragment Vn-α-syn or in complex with α-syn-Vc). In a further study, Kiechle *et al.* investigated age-dependent changes to α-syn oligomerization and aggregation at the synapse in two transgenic mouse models where α-syn was fused to the N- or C-terminal part of hGaussia Luciferase or YFP Venus halves (Kiechle et al., 2019). For the hGaussia Luciferase protein-fragment system, the two complementation fragments were expressed at different levels but for the Venus system, the author presented data that showed the two BiFC fragments both equally and differentially expressed. In the cases when expression was differential, findings were consistent with ours as levels of α-syn-Vc were reduced.

A possible explanation for the difference in protein levels could be due to the arrangement of the BiFC fusion proteins (Fig. 1); the split Venus fragments fused at the opposite ends of α-syn may influence α-syn stability or clearance. Similar to α-syn, differences in protein expression levels have also been consistently observed when Venus-BiFC has been fused with, amyloid precursor protein (APP) (So et al., 2013) and exon1 of the huntingtin protein (Httex1) (Herrera et al., 2011) but not in the Luciferase- or GFP-based BiFC systems when both complementation constituents are fused to the carboxy-terminus of α-syn (Mohammadi et al., 2017; Moussaud et al., 2015; Outeiro et al., 2008). The large differences in the protein levels of the two Venus BiFC fragments combined with the high aggregation propensity of Vn-α-syn highlight the critical importance of assessing the expression and aggregation state of each component in the Venus BiFC systems as well as other BiFC systems to ensure accurate interpretation of experimental findings. In the absence of such insight, it is difficult to correlate the differences in the fluorescence signal to the propensity of α-syn to oligomerize or speculate about the mechanisms of α-syn oligomerization, aggregation and toxicity in these cellular assays. This is especially important when the fluorescence signal arises from only a small fraction of the total α-syn expressed in the cell. Finally, whether the oligomers formed by Vn-α-syn could influence α-syn cellular properties in cellular assays using Venus-BiFC system remains unknown (Kiechle et al., 2019; Zondler et al., 2014). Together, these observations underscore the limitations of relying solely on the fluorescence signal as the primary readout of α-syn oligomerization.

Although previous studies did not thoroughly investigate the oligomerization properties of the individual BiFC fragments, two studies showed by denaturing (Roberts et al., 2015) and native gel electrophoresis (Outeiro et al., 2008; Roberts et al., 2015) that some GFP-BiFC fragments have the propensity to form high molecular weight oligomers on their own (Kerppola, 2008). Despite the fact that in both these studies and a previous report (Magliery et al., 2005) the fragment with GFP/Venus fused to the N-terminus of α-syn exhibited increased oligomer formation, the significance and implications of these observations were not discussed or accounted for in the interpretation of the results. In another study by Dettmer and colleagues, intact cell crosslinking followed by WB analysis showed the presence HMW species corresponding to Vn-α-syn only, which is again consistent with our findings (Dettmer et al., 2015). Furthermore, Bae *et al*. showed that only Vn-α-syn was secreted by cells and existed as a mixture of monomers and aggregates in the cell culture medium (Bae et al., 2014), consistent with our data (Fig. 4A). When the cell culture media containing Vn-α-syn monomers and aggregates were added to cells stably expressing α-syn-Vc, fluorescent foci were observed in the host cells. If the Vn-α-syn aggregates were removed prior to addition to other host cells, no BiFC signal was observed. These observations suggest that the fluorescence signal observed in the host cell arise due to the recruitment and/or entrapment of the α-syn-Vc monomers in the preformed Vn-α-syn aggregates rather than interactions between the two monomers, consistent with our observations and those of previous reports (Eckermann et al., 2015). This experiment illustrates the challenges in interpreting BiFC fluorescence signal in systems where different BiFC fragments exhibit differences in protein levels, biophysical properties and aggregation propensity and demonstrate that other aberrant interactions, in addition to the expected multimerization may give rise to the BiFC fluorescence signal.

One way to validate the cellular BiFC-based oligomerization assays is to assess the specificity of the interactions by using mutant forms of the protein of interest that exhibit no, or a very low, propensity to self-associate or by using closely related proteins that do not interact with one another or self-oligomerize (Hu et al., 2002; Kudla & Bock, 2016). In these cases, one would expect no, or very little, fluorescent signal when compared to that of BiFC protein partners that do oligomerize. Indeed, this approach has already been applied by several groups to investigate the utility of the Venus BiFC system to compare the oligomerization properties of the different members of the α-syn family (Eckermann et al., 2015) and various α-syn disease-associated mutants (Dettmer et al., 2015; Lázaro et al., 2016, 2014). For example, Eckermann and colleagues reported no detectable differences in the Venus BiFC signal when α-, β- and γ-syn oligomerization were assessed in parallel, suggesting that they exhibit similar homodimerization properties (Eckermann et al., 2015). This is surprising, as β-syn is known to not aggregate, does not form inclusions in cells or the brain and has consistently been shown not to form fibrils under physiological conditions (Ghosh et al., 2015). Similarly, one would expect an increased BiFC signal when using proteins with disease-specific point mutations such as A53T, that promote α-syn aggregation. Although several α-syn PD-associated mutants have been shown to promote α-syn oligomerization and/or fibril formation *in vitro* (de Oliveira & Silva, 2019; Ruf et al., 2019), in cells (M. B. Fares et al., 2014; Khalaf et al., 2014; Lei, et al., 2019) and *in vivo* (Mbefo et al., 2015; Paumier et al., 2013), Lázaro et al. reported identical Venus BiFC signals from WT α-syn and the PD-linked mutants A30P, E46K, H50Q, G51D and A53T (Lázaro et al., 2014). Similar observation were made by Outeiro et al using a GFP-BiFC cellular assay (Outeiro et al., 2008). It was suggested that these observations could be explained by the possibility that the dimerization propensity of the α-syn proteins to form dimers does not reflect their oligomerization and aggregation properties (Eckermann et al., 2015). However, this interpretation is not consistent with extensive biophysical studies using alternative approaches that show significant differences in the propensity of α-syn with the various PD-linked α-syn mutants to form oligomers (Winner et al., 2011) and dimers (Janowska et al., 2015; Lv et al., 2016). In addition, several studies have shown significant differences in the lifetime and distribution of different types of dimers formed by wild-type α-syn and PD-linked mutants (Krasnoslobodtsev et al., 2013; Krishnan et al., 2003; Lv et al., 2015). Taken together, these results demonstrate that the current α-syn BiFC cellular assays are not predictive of the propensity for α-syn to form oligomers or fibrils. Fundamentally, these data highlight that the α-syn BiFC assay is limited to specific protein conformations and many of these are not visible during detection due to the nature of bifluorescent complementation. If using this in a drug discovery setting to identify or investigate inhibitors or modifiers of α-syn dimerization/oligomerization there is a high risk that active compounds may present as false negatives, which would likely result in poor translation of drug activity in other preclinical models.

Despite these limitations, several studies have demonstrated that the α-syn BiFC system could be useful to investigate α-syn cell-to-cell transmission (Bae et al., 2014; K. M. Danzer et al., 2011; Kim et al., 2016; Wang et al., 2014). In these experiments, each BiFC fragment is expressed in different cells which are then co-cultured and α-syn cell-to-cell transfer is monitored by the reconstitution of the BiFC fluorescence signal in either cell. In such studies, it is again crucial to biochemically assess the aggregation state of the BiFC fragment that is formed and secreted by each cell lines. These recommendations are in line with those made previously by Eckermann et al. (Eckermann et al., 2015).

## Conclusions

The design of BiFC systems to investigate protein-protein interactions or protein self-assembly is based on three assumptions 1) the individual reporter fragments do not self-associate unless the proteins they are fused to interact with each other or have the propensity to dimerize or oligomerize (Kerppola, 2006); 2) both BiFC fragments are expressed at equal levels; and 3) mutants that disrupt or enhance protein-protein interactions should result in a decrease or an increase in the fluorescence signal, respectively. While our studies confirm that a fluorescence signal is observed only when both BiFC fragments fused to α-syn are co-expressed in the cell, they also clearly demonstrate that 1) the Vn-α-syn and α-syn-Vc BiFC fragments are not expressed at equal levels; 2) the Vn-α-syn fragment exhibits a high propensity to aggregate; and 3) the fluorescence signal observed may not arise only from α-syn dimerization and could also be derived from the association or entrapment of α-syn-Vc in the Vn-α-syn aggregates. Together, these observations combined with previous findings from independent groups demonstrate that the α-syn-Venus-BiFC system does not reveal known differences in the oligomerization and aggregation propensities of PD-associated variants or the interactions between mutants and WT α-syn (Eckermann et al., 2015). Finally, consistent with previous reports, we showed that overexpression of either Vn-α-syn or the two BiFC fragments results in the formation of α-syn oligomers that do not seem to convert to inclusions or amyloid-like aggregates (Dettmer et al., 2015; Gonçalves et al., 2016; Kiechle et al., 2019; Lázaro et al., 2014; Moussaud et al., 2015; Outeiro et al., 2008). Together, these findings underscore the critical importance of characterizing BiFC-based cellular assays at the biochemical and structural levels and understanding how the fusion of fragments of fluorescent proteins influences α-syn biophysical and cellular properties before using such assays in mechanistic, target validation and drug discovery studies.

## Abbreviations

α-syn: α-synuclein
α-syn-V: α-synuclein-full length Venus
Vn-α-syn: n-terminal fragment of Venus fused to the amino-terminus of α-syn
α-syn-Vc: c-terminal fragment of Venus fused to the carboxy-terminus of α-syn
BiFC: bimolecular fluorescent complementation
PD: Parkinson’s disease
SEC: size exclusion chromatography

## Acknowledgments

This work was supported by funding from EPFL and by core institutional funding from Alzheimer’s Research UK. We are grateful to the Bioimaging Core Facility (BioP, EPFL) for their technical support. We would also like to thank Dr. Senthil Kumar for his technical help with size exclusion chromatography, Muhammed Syed for providing some of the recombinant proteins used as standards and in the SEC studies. Dr. Tim Newton assisted in the design of bidirectional plasmid and BacMam constructs. Dr. Anne-Laure Mahul for helpful discussions and Dr. Kolla Rajasekhar for generating the graphical abstract. James Duce is an editor for Journal of Neurochemistry. We would also like to thank Prof. Tiago Outeiro and Prof. Pamela McLean for providing us with the α-syn-Venus-BiFC expression vectors used in this study and Prof. Seung-Jae Lee and Eun-Jin Bae for providing the BiFC expressing cell lines, SH-SY5Y (ECACC) neuroblastomas stably expressing either Vn-α-syn or α-syn-Vc. The manuscript was published on BioRxiV before: https://www.biorxiv.org/content/10.1101/2020.05.02.074161v2.

---

**Supplementary Figure 1:**
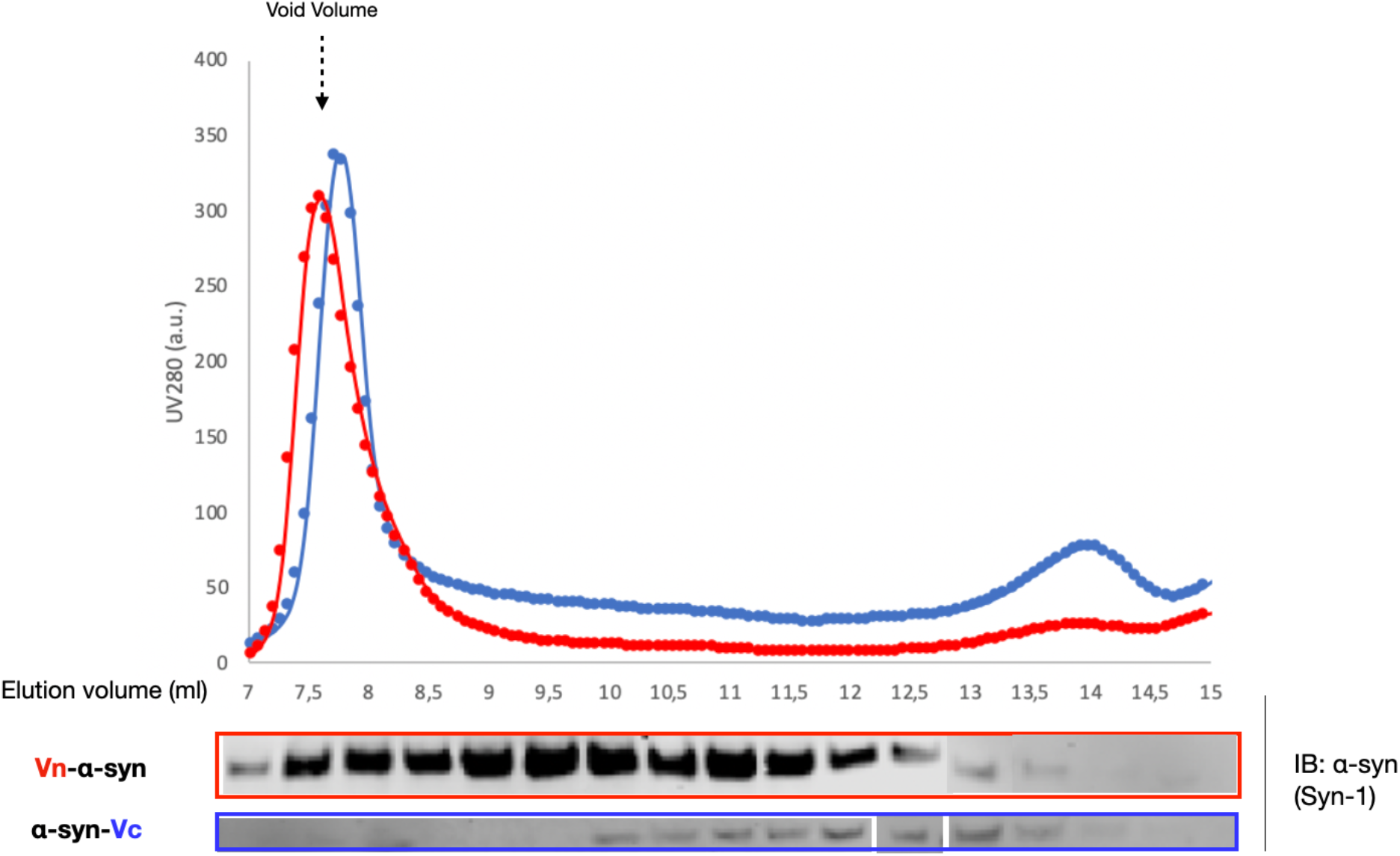
The Vn-syn fragment exhibits a high propensity to form soluble oligomers compared to α-syn-Vc and WT α-syn also in SH-SY5Y cells stably expressing these constructs. Size exclusion chromatography of SH-SY5Y cells that stably expressed the indicated constructs, were lysed 72h after plating. The SEC fractions were analyzed by WB using anti-α-syn (Syn-1) antibodies. Representative data shown from n=2 experiments. Vo: dead volume; Ve: elution volume; Vn-α-syn: (amino terminal fraction of Venus)-syn; α-syn-Vc: α-syn-(carboxy terminal fraction of Venus); α-syn-V (-synuclein-Venus full length).

**Supplementary Figure 2:**
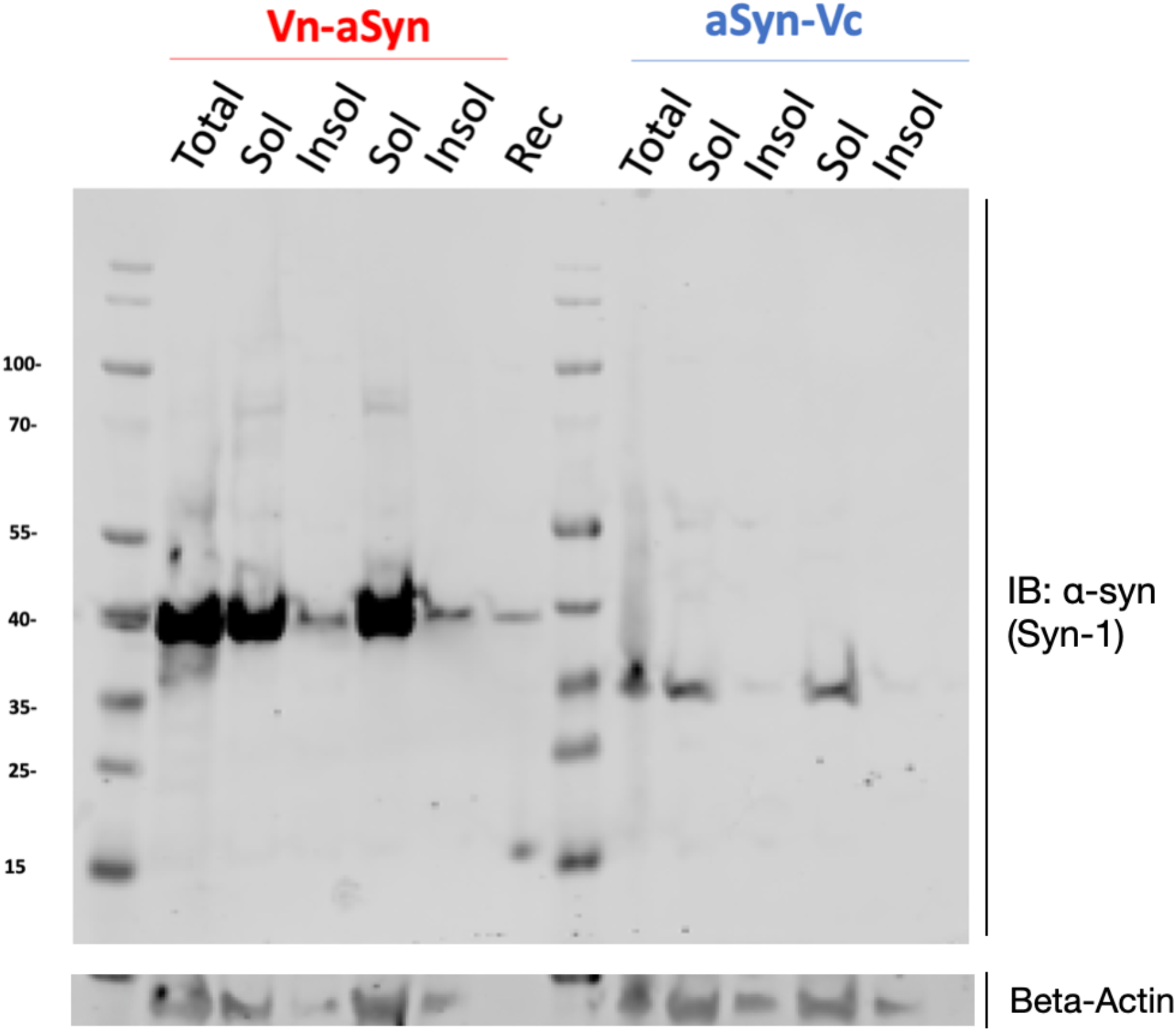
Vn-α-syn, but not α-syn-Vc or WT α-syn, is detected in the insoluble fractions. SH-SY5Y cells stably expressing the indicated α-syn constructs were lysed 72 h after plating, and the cell lysates were separated into a soluble and insoluble fraction by centrifugation. The resulting fractions were analyzed by immunoblotting of α-syn (Syn-1). Beta-Actin was used as the loading control. Shown is a representative blot of n=2 experiments. Vn-α-syn: (amino terminal fraction of Venus)-syn; α-syn-Vc: α-syn-(carboxy terminal fraction of Venus).

**Supplementary Figure 3:**
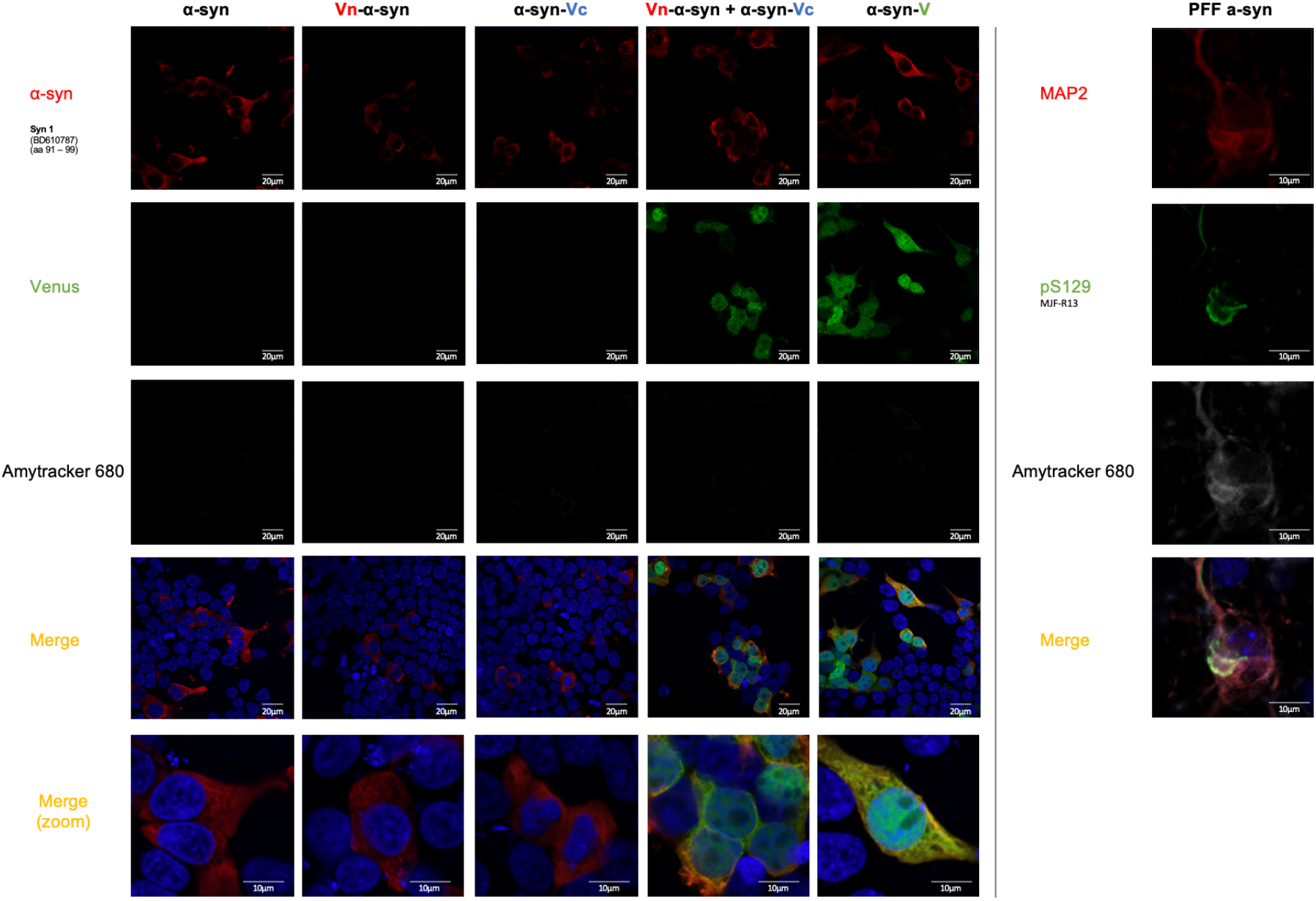
No amytracker positive inclusions were observed in cell lines expressing the BiFC fragments. HEK-293 cells expressing the indicated α-syn constructs were fixed 48 h post-transfection and stained with the Syn-1 antibody. Upon co-transfection of the Vn-α-syn and α-syn-Vc constructs, Venus was successfully reconstituted and emitted a fluorescent signal that colocalized with the signal for α-syn. Mouse primary neurons were treated by recombinant pre-formed-fibrils α-syn for 14 days and are positive for the amytracker. The nucleus was counterstained with DAPI. MAP2 was used as a neuronal marker for the primary neurons and also colocalize with the amytracker. Both α-syn-V and the reconstituted BiFC fragments showed a nuclear localization and signal that did not colocalize with that of α-syn (Syn-1) The scale bar is 20 μm in all the non-zoomed images and 10 μm for the zoomed ones. Shown are representative images from two independent experiments. The microscopy settings were kept the same among each construct.

## Notes

### Competing Interest Statement

The authors have declared no competing interest.

